# Association of Striatal Connectivity Gradients to Functional Domains Across Psychiatric Disorders

**DOI:** 10.1101/2022.06.02.494510

**Authors:** Peter C.R. Mulders, Philip F.P. van Eijndhoven, Jasper van Oort, Marianne Oldehinkel, Fleur A. Duyser, Josina D. Kist, Rose M. Collard, Janna N. Vrijsen, Koen V. Haak, Christian F. Beckmann, Indira Tendolkar, Andre F. Marquand

## Abstract

**Objective:** To uncover transdiagnostic domains of functioning across stress- and neurodevelopmental disorders, and to map these on to the topographic functional organization of cortico-striatal circuitry.

**Methods:** In a clinical sample (n=186) of subjects with high rates of comorbidity of major depressive disorder, anxiety disorder, attention-deficit/hyperactivity disorder and/or autism spectrum disorder, we use exploratory factor analysis on a wide range of clinical questionnaires to identify consistent functional domains of symptomatology across disorders, then replicate these functional domains in an independent dataset (n=188). Then, we use canonical correlation analysis link these functional domains to the topographic organization of the striatum as represented by connectopic maps.

**Results:** We reveal four functional domains that transcend current diagnostic categories relating to negative valence, cognition, social functioning and inhibition/arousal. These functional domains are replicated in an independent sample and are associated with the fine-grained topographical organization of functional connectivity in the striatum (out of sample r=0.20, *p*=0.026), a central hub in motor, cognitive, affective and reward-related brain circuits.

**Conclusions and relevance:** Functional domains across stress- and neurodevelopmental disorders are associated with the functional organization of the striatum. We propose that investigating psychiatric symptoms across disorders is a promising path to linking them to underlying biology, and can help bridge the gap between neuroscience and clinical psychiatry.

## Introduction

Psychiatric disorders are behaviorally and biologically complex, as is evidenced by our syndrome-level understanding of their clinical phenomenology. While a common framework is critical for providing patient care and evaluating the efficacy of therapeutic options, classification systems that are commonly used to define psychiatric disorders also constrain the way in which we are able to connect these disorders to underlying biology (1). Moreover, these symptom-based diagnostic systems typically reflect a convergence of multiple distinct biological mechanisms, contributing to significant clinical heterogeneity within disorders and high levels of comorbidity between them (2, 3). This comorbidity can also stem from a single biological origin driving various symptoms that are classified separately, which may help explain treatment being effective for different classified disorders. Our underlying neurobiology is not constrained to classification paradigms, which is exemplified by various brain regions, genes and neurobiological pathways being implicated across the psychiatric spectrum (4, 5). This lack of distinct underlying biological features for psychiatric disorders has impaired clinical progress, for example in the search for reliable neuroimaging biomarkers for the presence or prognosis of mental disorders (6). The MIND-Set (“Measuring Integrated Novel Dimensions in Neurodevelopmental and Stress-related Mental Disorders”) cohort was initiated with this in mind, employing concepts from transdiagnostic groundwork such as the Research Domain Criteria, to collect data across neurodevelopmental and stress-related disorders and investigate their clinical and biological (co)variation (7). These transdiagnostic frameworks focus on domains of functioning across diagnoses that could prove valuable from a research perspective, as well as from the perspective of understanding patient functioning at the individual level (1).

Within such frameworks, neuroimaging studies have made significant progress in linking specific domains of functioning to brain networks and regions across disorders (6, 8, 9). These studies suggest several key regions and circuits being relevant in the majority of psychiatric disorders, with related disorders (e.g. mood and anxiety disorders) showing strongest similarities with one another in terms of their association with distributed patterns of brain function and/or structure. The striatum, which has been implicated across the behavioral spectrum, is such a key locus of convergence for psychopathology across multiple disorders. As a central hub in a range of motor, cognitive, affective and reward-related brain circuits, the striatum receives a large array of lateral and medial cortical inputs, which are topographically organized (10, 11). In addition, it receives afferents from the thalamus and dopaminergic input from the brain stem (12), while the striatum itself projects mainly to other basal ganglia. Via extensive connectivity with the (pre)frontal cortex, the striatum is critical in learning, adapting and executing goal-directed behavior, taking into account the complexity that leads to behavior such as emotion and cognition. Being involved in the entire spectrum of goal-directed behavior from sensory through motivational to cognitive and executive function, it is unsurprising that changes in striatal structure or function have been extensively reported across a wide range of psychiatric disorders (13–18). In addition, inter-individual differences in striatal structure and function have been shown to reflect disease severity across psychotic disorders, depressive disorders, post-traumatic stress disorder and obsessive-compulsive disorder, regardless of which specific disorder was present (19). These factors make the striatum a target of significant interest when investigating broad functional domains impaired across psychiatric disorders.

The striatum is extensively connected with almost the entire cortex and is involved in multiple behaviorally relevant circuits which can be probed, for example, through functional connectivity. However, connectivity in the striatum is topographically organized, such that nearby regions in the striatum are connected with nearby regions in cortex and therefore conventional functional connectivity approaches are unable to capture the complexity of functional connections within this complex system, nor reflect the topographic characteristics of striatal organization at the single-subject level. Through recent advancements, we can now investigate the topographic organization of striatal circuits through connectopic maps (20). These maps, which are estimated at the single-subject level, represent slowly varying topographically organized connectivity patterns (‘connectopic gradients’) that reveal how connectivity to the rest of the brain is organized within a region of interest, even when multiple connectivity patterns are spatially overlapping but functionally distinct. The latter is important because in striatum we have shown that the dominant mode of connectivity change distinguishes between caudate nucleus, nucleus accumbens and putamen, while the second mode follows a rostral-caudal gradient across the three striatal substructures (21). We have also shown that topographic connectivity in the striatum is related to complex goal-directed behaviors at the individual level (21), and shows a strong correspondence with the spatial distribution of dopaminergic projections, demonstrating their potential for investigating striatal function (22). Because connectopic maps characterize complex regions such as the striatum in a way that links to behavior, we hypothesize that individual differences in connectopic maps are predictive of psychiatric symptomatology across disorders.

In this study, we apply connectopic mapping to a richly phenotyped, transdiagnostic and highly comorbid cohort – the ‘Measuring Integrated Novel Dimensions in neurodevelopmental and stress-related mental disorders (MIND-Set) study – aiming to (i) dissect psychiatric symptom profiles within a heterogeneous and clinically representative sample, providing quantitative measures across different domains of functioning that cut across diagnoses and measurement instruments and (ii) show that these domains of functioning are linked to the topographic functional organization of the striatum at the level of the individual patient. To achieve these goals, we apply an innovative multivariate analytical strategy that combines penalized canonical correlation analysis with stability selection that provides the ability to learn brain-behavior associations whilst providing unbiased estimates of generalizability and strong statistical guarantees over false detection of associations.

## Methods

A detailed description of analytical procedures is provided in the supplement. In brief, data were collected as part of the MIND-Set study, an observational, naturalistic, cross-sectional study that includes adult patients with stress-related and/or neurodevelopmental disorders that were assessed at the outpatient unit of the department of psychiatry at the Radboud university medical center (Radboudumc) in Nijmegen, the Netherlands. For a more detailed description of the study design and set-up we refer to previous work (7). Importantly, individuals in this sample have high levels of comorbidity, accurately reflecting the clinical reality.

### Study participants

*Discovery/neuroimaging sample*: participants from the MIND-Set cohort were included who met criteria for at least one of the following psychiatric disorders: major depressive disorder, anxiety disorder, attention-deficit/hyperactivity disorder (ADHD) and autism spectrum disorder (ASD), and had completed behavioral and neuroimaging assessments (*n* = 203, before connectopic mapping). Diagnoses were confirmed using the Structured Clinical Interview for DSM-IV-TR (SCID-I/P) (23), the Diagnostic Interview for Adult ADHD (DIVA) (24) and/or the Nijmegen Interview for Diagnosing adult Autism spectrum disorders (NIDA) (25). *Replication sample*: n=188 (101 men / 87 women; age 43.6±14.3 years; MDD=61, anxiety=54, ADHD=54, ASD=38). This replication sample followed the same inclusion process and deep phenotyping as the main sample, with the exception of an MRI session.

### Behavioral data and factor analysis

We used an extensive panel of questionnaires covering multiple symptom and functional domains (see table 2). Exploratory factor analysis (EFA) was used to uncover four domains of functioning that transcend conventional diagnostic (DSM) boundaries.

### Connectopic mapping and statistical analysis

Imaging data was processed using standard pipelines with FSL 5.0.11 (FMRIB, Oxford, UK) and careful attention was given to addressing known confounds in resting FMRI, such as head motion (see supplementary data) (26, 27). We applied *ConGrads*, a fully data-driven method based on manifold learning and spatial statistics, to the resting-state fMRI data to obtain highly individualized representations of striatal functioning (‘connectopic maps’) for each subject (20). These maps represent slowly varying topographic patterns of connectivity (‘gradients’) that map connectivity changes within a target region in relation to the rest of the brain at the individual subject level. Although we focused on the principal or dominant gradient, multiple overlapping topographic representations can exist simultaneously within a single region, so both principal and second gradients were estimated to be able to investigate other potential effects driving associations to behavior. All gradients were visually inspected, and subjects were excluded if a clear gradient could not be estimated, due to imaging artefacts or spatial correlation of individual gradients to the group maps was low (n=17). For each subject, trend surface coefficients fit to the gradients from each hemisphere were used to provide a low-dimensional summary of the connectopy, following prior work (20, 21). These were concatenated for each subject and entered into a penalized canonical correlation analysis (CCA) model (28). The CCA was used to determine the association between the behavioral domains of functioning and striatal gradients, using feature stability selection (29) and out-of-sample testing. This procedure has been described in detail in earlier publications (28), is summarized in figure 1, and detailed in the supplementary information. To establish model consistency, we performed the full analysis ten times using different (stratified) data splits. Finally, we calculated the (cross-)loadings of brain features to the different functional domains to understand the relative contribution of different behavioral and neurobiological features to the associations.

**Figure 1.**
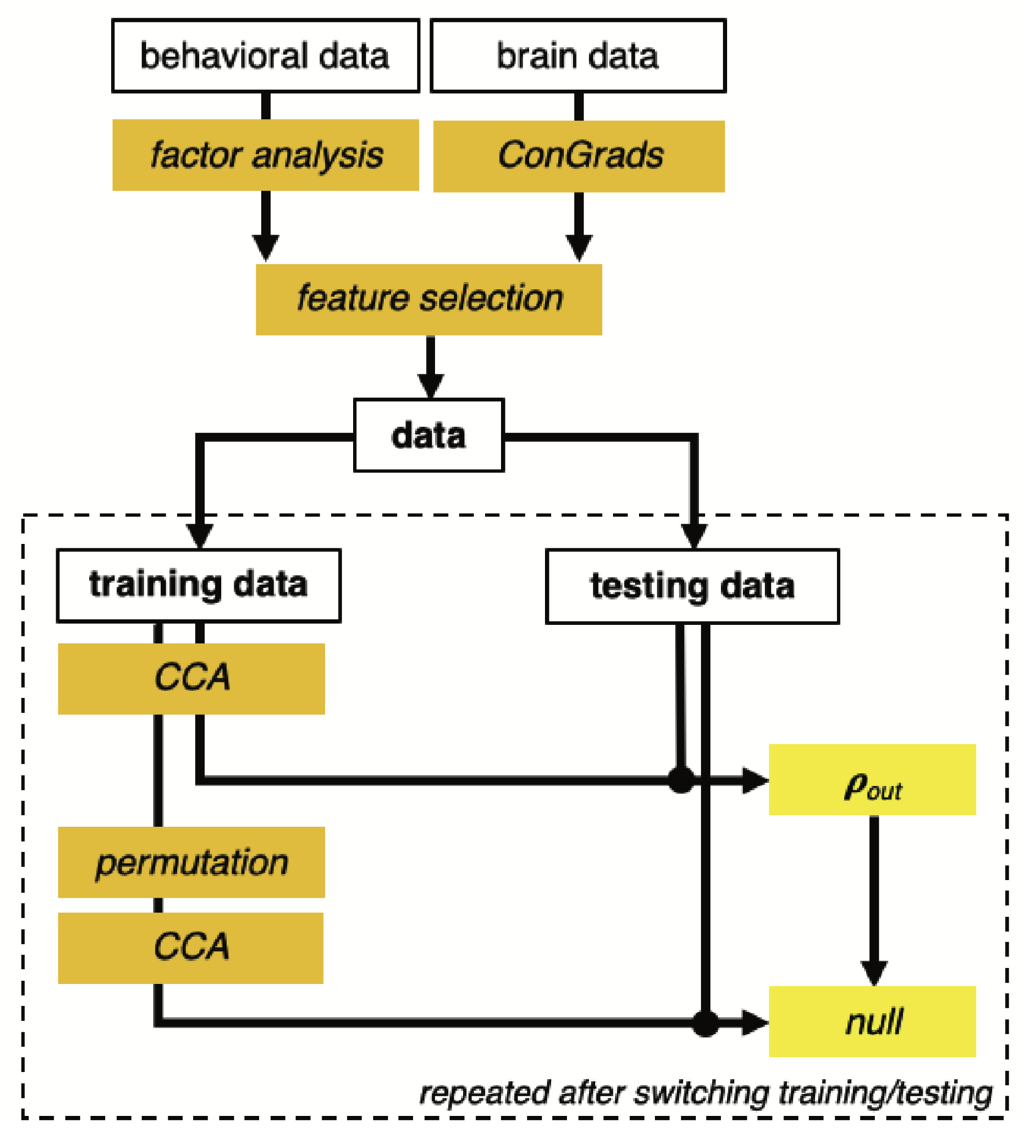
Processing and analysis pipeline. Exploratory factor analysis is used to obtain functional domain scores, while ConGrads is used to obtain functional gradient trendsurface coefficients. Data were split (50/50), and feature stability selection was performed on the training data to ensure stability. CCA was used to obtain the canonical correlates and weights, which were used on the training data to obtain out-of-sample canonical correlates and correlations. Data was permuted (500 permutations) to obtain the corresponding null. CCA and permutation were repeated after switching training and testing data. The full analysis was performed 10 times to reduce the risk of type 1 errors.

## Results

A total of 186 subjects were included in the final analysis (107 men, 79 women; age 37.9±13.2 years). As per the study design, the sample contained high levels of comorbidity across symptoms and diagnoses: 102 subjects had a single diagnosis, 69 had two diagnoses, 13 had 3 diagnoses and 2 subjects had all included disorders (table 1). Variation in comorbidity is presented in figure 2a.

**Table 1.** Demographics and questionnaire data for full sample and per diagnostic category. Note that subjects can be in multiple categories. Abbreviations: Inventory of Depressive Symptomatology (IDS), Anxiety Sensitivity Index (ASI), Connor’s Adult ADHD Rating Scale (CAARS), Autism-Spectrum Quotient (AQ-50), Brief Assessment of Impaired Cognition Questionnaire (BRIEF), Personality Inventory for DSM-IV Short Form (PID), Toronto Alexithymia Scale (TAS-20) and the Perseverative Thinking Questionnaire (PTQ).

|  | full | depression | anxiety | ADHD | ASD |
| --- | --- | --- | --- | --- | --- |
| *n* | 186 | 89 | 59 | 73 | 66 |
| *gender (m/f)* | 107 / 79 | 49 / 40 | 28 / 31 | 45 / 28 | 44 / 22 |
| *age (y)* | 37.9 ± 13.2 | 40.1 ± 14.3 | 36.4 ± 12.4 | 34.4 ± 10.4 | 35.3 ± 12.5 |
| *IDS* | 33 ± 13 | 41 ± 11 | 35 ± 12 | 30 ± 13 | 30 ± 12 |
| *ASI* | 15 ± 9 | 17 ± 10 | 18 ± 9 | 14 ± 8 | 15 ± 9 |
| *CAARS* | 18 ± 6 | 19 ± 6 | 19 ± 5 | 20 ± 5 | 18 ± 5 |
| *AQ-50* | 126 ± 20 | 124 ± 21 | 132 ± 17 | 122 ± 18 | 138 ± 17 |
| *BRIEF* | 142 ± 22 | 144 ± 22 | 143 ± 19 | 149 ± 19 | 139 ± 23 |
| *PID* | 28 ± 10 | 29 ± 10 | 30 ± 9 | 28 ± 10 | 29 ± 9 |
| *TAS-20* | 56 ± 12 | 57 ± 12 | 58 ± 9 | 53 ± 12 | 58 ± 11 |
| *PTQ* | 36 ± 9 | 38 ± 10 | 38 ± 8 | 34 ± 9 | 37 ± 9 |

**Figure 2.**
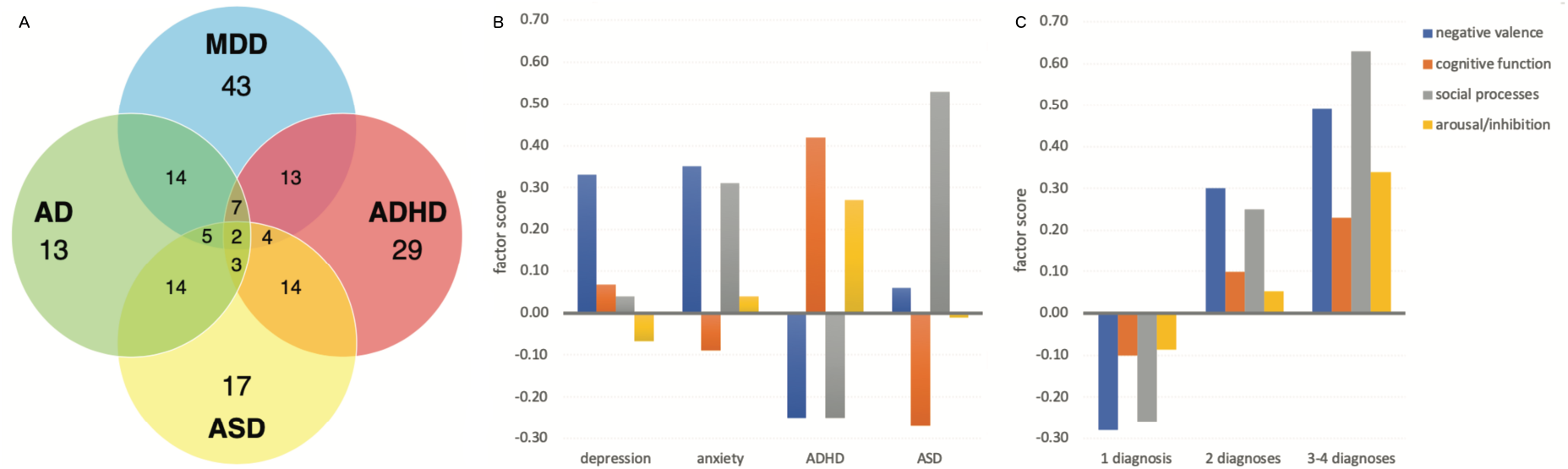
Comorbidity and functional domains. **A**: Diagram showing comorbidity within the data. **B:** Average subject loading onto the four functional domains in relation to diagnosis **C**: average subject loading onto functional domains in relation to degree of comorbidity. MDD: major depressive disorder, AD: anxiety disorder, ASD: autism spectrum disorder, ADHD: attention-deficit hyperactivity disorder.

### Factor analysis and functional domains

Exploratory factor analysis on 31 scales and subscales across psychiatric symptomatology decomposed behavioral data into four factors (Kaiser-Meyer-Olkin test 0.85, see table 2). The first factor relates to negative valence systems (1) and contains scores relating to depressive and anxiety symptoms, in addition to negative affect and repetitive negative thinking scores. The second factor includes symptoms related to cognitive function, including planning, organizing, working memory and mental capacity. The third factor relates to social functioning and includes social awareness, communication, and external awareness. The fourth factor includes symptoms relating to (dis)inhibition, antagonism and emotional lability. Broadly, we interpret these four factors as representing four functional domains: 1) negative valence, 2) cognitive function, 3) social processes and 4) arousal/inhibition. These functional domains transcend diagnostic boundaries and each factor includes scales and subscales from questionnaires probing different underlying disorders. As expected, these factors were not independent but were correlated to one another. Factor analysis on the replication sample of 188 subjects showed almost identical grouping of behavioral data (table 2; correlation of factor loadings r = 0.90, p < 0.001).

**Table 2.**
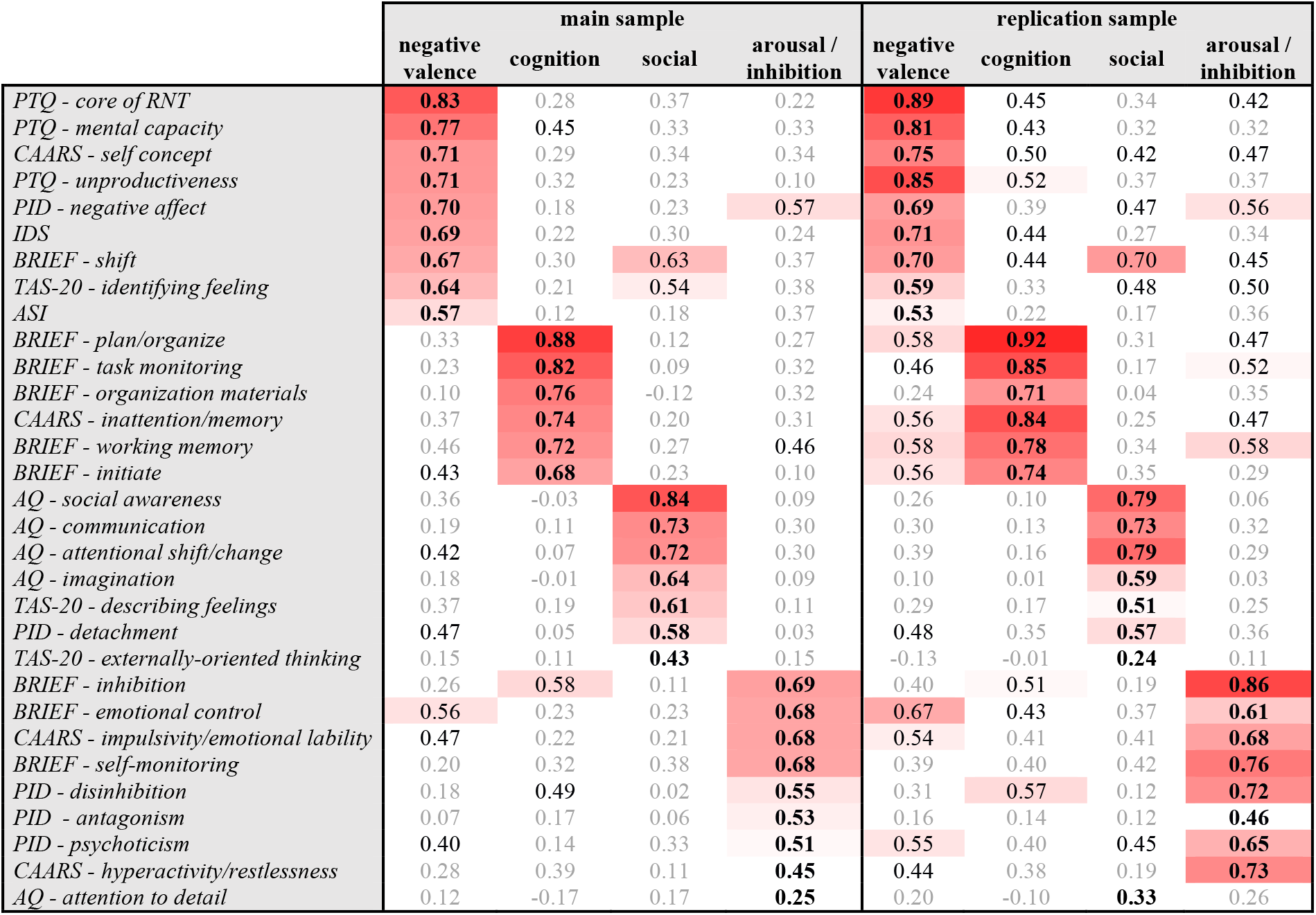
Results of exploratory factor analysis and factor loadings. Color intensity relates to strength of the behavioral item within the functional domain (>0.5). Bold indicates the highest factor loading for the behavioral item. Abbreviations: RNT: Repetitive negative thinking; AQ-50: Autism-Spectrum Quotient; CAARS: Connor’s Adult ADHD Rating Scale; IDS: Inventory of Depressive Symptomatology; ASI: Anxiety Sensitivity Index; PID: Personality Inventory for DSM-IV Short Form; BRIEF: Brief Assessment of Impaired Cognition Questionnaire; TAS-20: Toronto Alexithymia Scale; PTQ: Perseverative Thinking Questionnaire.

|  | main sample |  |  |  | replication sample |  |  |  |
| --- | --- | --- | --- | --- | --- | --- | --- | --- |
|  | negative valence | cognition | social | arousal / inhibition | negative valence | cognition | social | arousal / inhibition |
| PTQ - core of RNT | **0.83** | 0.28 | 0.37 | 0.22 | **0.89** | 0.45 | 0.34 | 0.42 |
| PTQ - mental capacity | **0.77** | 0.45 | 0.33 | 0.33 | **0.81** | 0.43 | 0.32 | 0.32 |
| CAARS - self concept | **0.71** | 0.29 | 0.34 | 0.34 | **0.75** | 0.50 | 0.42 | 0.47 |
| PTQ - unproductiveness | **0.71** | 0.32 | 0.23 | 0.10 | **0.85** | 0.52 | 0.37 | 0.37 |
| PID - negative affect | **0.70** | 0.18 | 0.23 | **0.57** | **0.69** | 0.39 | 0.47 | **0.56** |
| IDS | **0.69** | 0.22 | 0.30 | 0.24 | **0.71** | 0.44 | 0.27 | 0.34 |
| BRIEF - shift | **0.67** | 0.30 | **0.63** | 0.37 | **0.70** | 0.44 | **0.70** | 0.45 |
| TAS-20 - identifying feeling | **0.64** | 0.21 | **0.54** | 0.38 | **0.59** | 0.33 | 0.48 | 0.50 |
| ASI | **0.57** | 0.12 | 0.18 | 0.37 | **0.53** | 0.22 | 0.17 | 0.36 |
| BRIEF - plan/organize | 0.33 | **0.88** | 0.12 | 0.27 | **0.58** | **0.92** | 0.31 | 0.47 |
| BRIEF - task monitoring | 0.23 | **0.82** | 0.09 | 0.32 | 0.46 | **0.85** | 0.17 | **0.52** |
| BRIEF - organization materials | 0.10 | **0.76** | -0.12 | 0.32 | 0.24 | **0.71** | 0.04 | 0.35 |
| CAARS - inattention/memory | 0.37 | **0.74** | 0.20 | 0.31 | 0.56 | **0.84** | 0.25 | 0.47 |
| BRIEF - working memory | 0.46 | **0.72** | 0.27 | 0.46 | 0.58 | **0.78** | 0.34 | **0.58** |
| BRIEF - initiate | 0.43 | **0.68** | 0.23 | 0.10 | 0.56 | **0.74** | 0.35 | 0.29 |
| AQ - social awareness | 0.36 | -0.03 | **0.84** | 0.09 | 0.26 | 0.10 | **0.79** | 0.06 |
| AQ - communication | 0.19 | 0.11 | **0.73** | 0.30 | 0.30 | 0.13 | **0.73** | 0.32 |
| AQ - attentional shift/change | 0.42 | 0.07 | **0.72** | 0.30 | 0.39 | 0.16 | **0.79** | 0.29 |
| AQ - imagination | 0.18 | -0.01 | **0.64** | 0.09 | 0.10 | 0.01 | **0.59** | 0.03 |
| TAS-20 - describing feelings | 0.37 | 0.19 | **0.61** | 0.11 | 0.29 | 0.17 | **0.51** | 0.25 |
| PID - detachment | 0.47 | 0.05 | **0.58** | 0.03 | 0.48 | 0.35 | **0.57** | 0.36 |
| TAS-20 - externally-oriented thinking | 0.15 | 0.11 | **0.43** | 0.15 | -0.13 | -0.01 | **0.24** | 0.11 |
| BRIEF - inhibition | 0.26 | **0.58** | 0.11 | **0.69** | 0.40 | 0.51 | 0.19 | **0.86** |
| BRIEF - emotional control | **0.56** | 0.23 | 0.23 | **0.68** | **0.67** | 0.43 | 0.37 | **0.61** |
| CAARS - impulsivity/emotional lability | 0.47 | 0.22 | 0.21 | **0.68** | 0.54 | 0.41 | 0.41 | **0.68** |
| BRIEF - self-monitoring | 0.20 | 0.32 | 0.38 | **0.68** | 0.39 | 0.40 | 0.42 | **0.76** |
| PID - disinhibition | 0.18 | **0.49** | 0.02 | **0.55** | 0.31 | **0.57** | 0.12 | **0.72** |
| PID - antagonism | 0.07 | 0.17 | 0.06 | **0.53** | 0.16 | 0.14 | 0.12 | **0.46** |
| PID - psychoticism | 0.40 | 0.14 | 0.33 | **0.51** | **0.55** | 0.40 | 0.45 | **0.65** |
| CAARS - hyperactivity/restlessness | 0.28 | 0.39 | 0.11 | **0.45** | 0.44 | 0.38 | 0.19 | **0.73** |
| AQ - attention to detail | 0.12 | -0.17 | 0.17 | **0.25** | 0.20 | -0.10 | **0.33** | 0.26 |

Regarding the distribution of factor coefficients within the different diagnostic classifications, we observed that while variance within groups was high, on average each diagnostic category showed a pattern that resembles its clinical manifestation (figure 2b). We also observed that as the degree of comorbidity increases (i.e. quantified by number of diagnoses for each individual), so did the loading onto all four domains of functioning (figure 2c).

### Connectopic mapping and canonical correlation analysis

At the group-level, striatal connectopic maps showed a best fit using up to a third order polynomial. The principal connectopic maps showed striatal topography similar to previous work (21), where it follows structural boundaries and gradually changes from putamen, to nucleus accumbens, to caudate (figure 3a). Projection maps for the striatal gradient onto the rest of the brain were also in line with previous work (figure 3b). As loading onto behavior was impacted by degree of comorbidity, we explored the impact of comorbidity on the connectopic maps and found that in highly comorbid subjects, the functional gradient was more pronounced (figure 3c). In other words, when observing the gradient in full there was a stronger separation between both ends of the functional spectrum, but the gradient direction was identical, possibly indicating more distinct connectivity patterns in subjects with high comorbidity.

**Figure 3.**
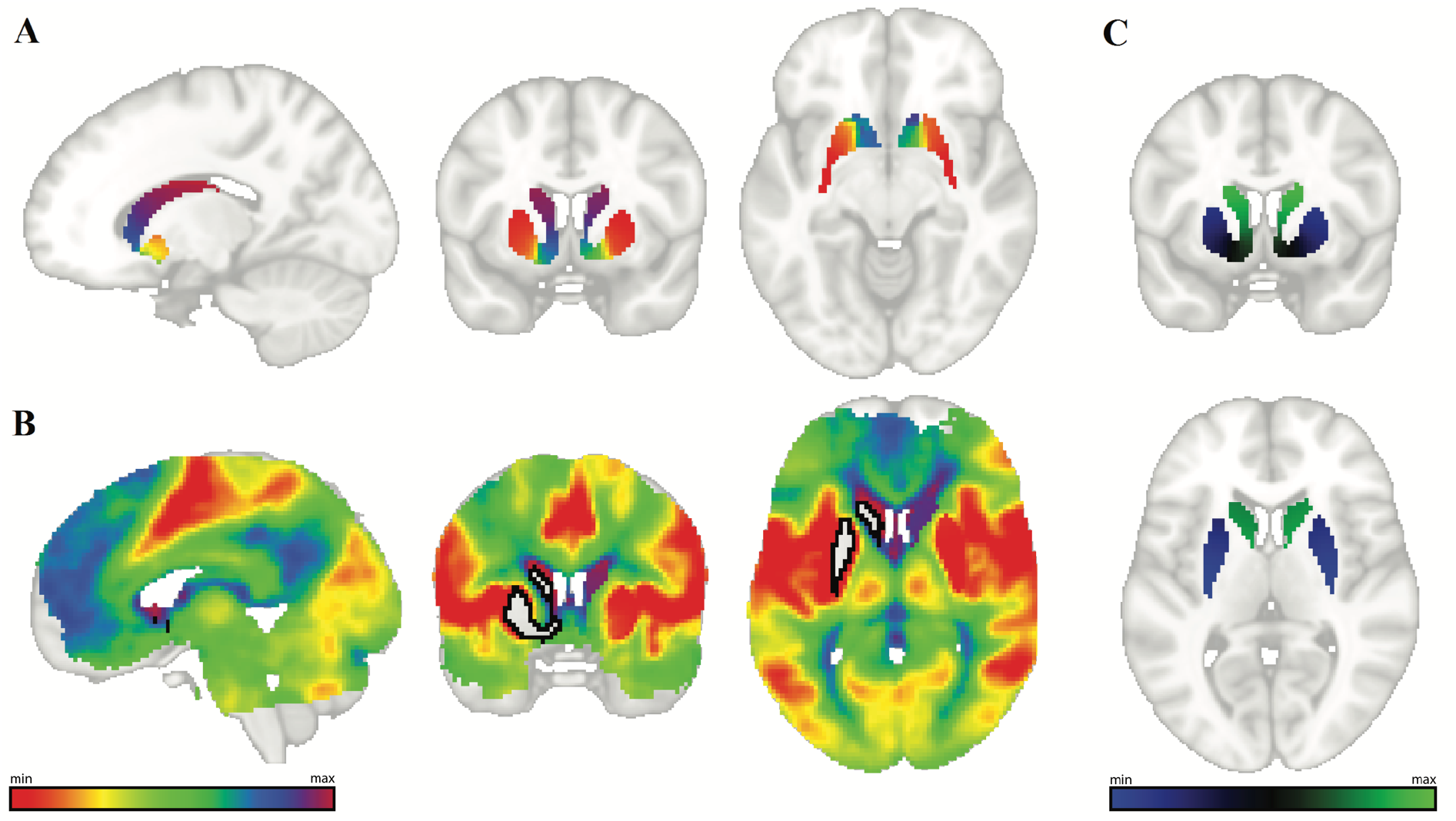
Results of connectopic mapping. **A**: average principal striatum gradient. **B**: average projection of left striatum gradient (black outline: left striatum), colors match those in A. **C**: average difference between subjects without comorbidity and with significant comorbidity (three or four diagnoses). The difference shown here indicates that while the gradient direction remains the same, with higher comorbidity the gradient becomes more pronounced, i.e. when observing the gradient in full there is a stronger functional separation between both ends of the functional spectrum.

Functional domain scores and trend surface coefficients were entered into a penalized canonical correlation analysis. Stability selection revealed the first and third-order trend surface coefficients in the *x*-direction of the left striatum as stable brain features. We found a clear and consistent correlation between functional domain scores and the connectivity gradients in the striatum (average in-sample correlation r=0.25±0.04, out-of-sample correlation r=0.20±0.02, *p*=0.026). This interaction explained variance across three of the four behavioral domains (negative valence 2.5%, cognitive 2.6%, social 0.2%, arousal/inhibition 3.6%), comparable in size to previous CCA-derived brain-behavioral associations (28). While these results were driven by left striatum organization within the model, no differences in gradients between left and right striatum were observed.

Additional tests were performed to determine whether our results could be explained by other effects (see supplementary information). Firstly, we showed that the reported association is specific to this particular topographic representation by repeating the full analysis using the second functional striatal gradient. Secondly, we showed that functional effects we report cannot be explained by underlying structural differences by repeating the analysis using classical volumetric measures of the striatum and finally, we showed that striatal topographic organization is not associated with the conventional diagnostic labels by repeating the CCA using the diagnostic labels instead of the functional domain scores. None of these other data modalities revealed any significant interactions between behavior and striatum structure or function.

## Discussion

In this work, we show how complex behavioral data in a sample with high psychiatric comorbidity can be represented in stable and reproducible domains of functioning. These functional domains are cross-diagnostic in that each of these domains incorporate parts from different questionnaires, and reproducible in that we were able to derive effectively identical latent structure across two independent samples. These domains of functioning are also similar to broad research areas in cognitive neuroscience in both healthy and clinical populations (1). We found one domain strongly tied to negative emotion and repetitive negative thinking (negative valence), one tied to organization and broad cognition, a domain related to social perception and functioning, and a domain related to arousal and (dis)inhibition. We observed that at the group level classified disorders showed a typical pattern of functional impairment across all four functional domains. The functional domains also showed a sensible grouping of symptoms that are known to be present within psychiatric patients, such as depressive symptoms co-occurring with repetitive negative thinking and problems in self-conceptualization. By grouping related symptomatology across disorders and across measurement instruments, these domains could be used to identify specific targets of treatment. As an example, repetitive negative thinking can be present in both depression and anxiety disorders and are grouped within the negative valence system. Treatments designed to affect negative valence systems dysfunction, such as cognitive behavioral therapy, might be effective for both disorders because they target this underlying functional domain.

We establish that these functional domains are related to the functional organization of the striatum across disorders. The multivariate nature of the CCA analysis approach employed means that changes in the overall gradient are associated with a pattern comprising multiple clinical measures. There is a large body of literature that supports our findings in linking the functional domains we observed to striatal function and connectivity. The ventral striatum is the key node within reward-circuitry, and crucial in the context of arousal and (dis)inhibition (11). Within the fronto-striatal circuit the striatum plays an important role in regulating behavior in response to salient stimuli. Differences in fronto-striatal circuitry have been at the forefront of neuroimaging research into disorders marked by strong disinhibition such as ADHD, oppositional defiant disorder, conduct disorder and addiction (30–32). The ventral striatum and the nucleus accumbens are also well established as important areas for affect (33), and our findings are in line with other studies that have found striatal connectivity changes in (remitted) depression and depressive symptoms (34, 35). With respect to the cognitive functional domain, the striatum and its dopaminergic modulation regulate those parts of cognition related to goal-directed behavior, such as working memory and attention switching (36). In fact, striatal markers based on neonatal imaging are predictive of cognitive ability years later (37). Changes in striatal morphology are also a distinguishing feature of neurodegenerative disorders marked by strong cognitive decline such as Alzheimer’s and Parkinson’s disease (38), disorders with high psychiatric comorbidity.

Taken together, known associations between striatal function and several functional domains in both clinical and healthy populations corroborate our findings. Furthermore, several studies have found striatal structure and/or function to be a marker for general symptoms, regardless of diagnosis (19). The nature of its impact on systems that are core to dysfunction in the context of psychiatric disorders explains limited disorder-specificity, while also underscoring the relevance of investigating functions over classifications. With regards to the underlying biology, we observed brain-behavioral interactions in the dominant functional gradient, where earlier work found interactions to behavior in higher-level gradients (22, 39). This indicates the complexity of the functional organization of the striatum, but also the necessity of investigating spatially overlapping patterns of connectivity in order to fully understand striatal involvement.

Several limitations to the current work need to be addressed. Firstly, the MIND-Set sample is highly comorbid by design, so replication and generalization studies in less complex or healthy samples are warranted to demonstrate transferability of the functional domains. Secondly, while substantial for a highly heterogeneous cohort, our sample size is moderate. Although balanced by careful assessment procedures including rigorous out-of-sample validation and stability selection (29), repeating the analysis in larger samples with a broader scope of biological targets could further uncover related circuitry.

In conclusion, we show how psychiatric symptomatology can be deconstructed into underlying functional domains that are reflected in striatal function. We believe that this transdiagnostic approach, which enables investigating domains of functioning that still carry the signature of the classified disorder while also incorporating individual variation that transcends the label, has great potential in overcoming current limitations in clinical and computational psychiatry. With a stronger biological footing and individualized nature, the functional domains could prove valuable in predicting clinical outcome. Through transdiagnostic research and understanding how disruptions in neural circuitry give rise to non-specific psychiatric symptoms or shared symptoms across different disorders, we will be able to pave the way for personalized treatment targeting circuits, regardless of which classified disorder is present.

## Acknowledgements

The authors also would like to thank professor Aart Schene for his continuous support in setting up the MIND-Set cohort. Without him, this work would not have been possible. Unfortunately professor Schene passed away too early to see the benefit of his continued support.

AFM gratefully acknowledges support from the European Research Council (ERC, grant ‘MENTALPRECISION’ 10100118).

