## supplemental information for "Association of Striatal Connectivity Gradients to Functional Domains Across Psychiatric Disorders"

### Supplementary information

#### Detailed methods and statistical analysis

Data were collected as part of the MIND-Set study ("Measuring Integrated Novel Dimensions in neurodevelopmental and stress-related mental disorders"), an observational, cross-sectional, naturalistic study that includes adult patients with stress-related and/or neurodevelopmental disorders that were assessed at the outpatient unit of the department of psychiatry at the Radboud university medical center (Radboudumc) in Nijmegen, the Netherlands. For a more detailed description of the study design and set-up we refer to previous work (1).

*Study participants*

Participants from the MIND-Set cohort were included if they met criteria for at least one of the following psychiatric disorders: major depressive disorder, anxiety disorder, attention-deficit/hyperactivity disorder (ADHD) and autism spectrum disorder (ASD), and had completed behavioral and neuroimaging assessments ($n$ = 203). Diagnoses were confirmed using the Structured Clinical Interview for DSM-IV-TR (SCID-I/P) (2), the Diagnostic Interview for Adult ADHD (DIVA) (3) and/or the Nijmegen Interview for Diagnosing adult Autism spectrum disorders (NIDA) (4).

*Behavioral data and factor analysis*

We used questionnaire data covering multiple symptom and functional domains, including Dutch versions of the Autism-Spectrum Quotient (AQ-50), Connor's Adult ADHD Rating Scale (CAARS), Inventory of Depressive Symptomatology (IDS), Anxiety Sensitivity Index (ASI), Personality Inventory for DSM-IV Short Form (PID), Brief Assessment of Impaired Cognition Questionnaire (BRIEF), Toronto Alexithymia Scale (TAS-20) and the Perseverative Thinking Questionnaire (PTQ). The 31 scores that were derived from these questionnaires and used in our analysis are listed in table 2. All these questionnaires are clinically used and validated, cover symptom dimensions of the included disorders and underlying pathology, and/or have been used in transdiagnostic research.

Exploratory factor analysis (EFA) in SPSS was used to uncover domains of functioning that transcend conventional diagnostic (DSM) boundaries. Parallel analysis (5) and scree-plots showed an optimal four-factor solution (maximum likelihood estimation, oblique rotation). To determine its robustness, EFA was repeated on an independent MIND-Set sample of similar size (n=188). This replication sample followed the same inclusion process and deep phenotyping as the main sample, with the exception of an MRI session.

*Magnetic resonance imaging and preprocessing*

Structural images (resolution 1.0 [$mm^3$]) were acquired using a T1-weighted Magnetization Prepared Rapid Acquisition Gradient Echo (MP-RAGE; TE/TR = 3.03/2300 [ms], flip angle = 8°, FOV = 256 × 256 × 192 [mm], GRAPPA acceleration factor 2). Resting-state images were collected using a 3T Siemens Magnetom Prisma MRI scanner (Erlangen, Germany) with a 32-channel head coil. T2*-weighted EPI BOLD-fMRI images were acquired for the resting-state scans, using a multi-band 6 protocol with an interleaved slice acquisition sequence (66 slices, TR = 1000 [ms], TE = 34 [ms], flip angle = 60°, voxel size 2.0 [$mm^3$], 500 volumes).

Preprocessing of resting state fMRI data was performed with FSL 5.0.11 (FMRIB, Oxford, UK) using the FMRI Expert Analysis Tool (FEAT) (6). The first five images were discarded, followed by brain extraction, motion and bias field correction, high-pass temporal filtering (100 s), and spatial smoothing with a 4 mm Gaussian kernel. Images were registered to standard space (MNI152) and ICA-AROMA was used for additional denoising, using 'non-aggressive' denoising as described in previous publications (7). Subjects were excluded if they showed more than 2 mm relative mean displacement.

*Connectopic mapping*
We applied *ConGrads*, a fully data-driven method using connectivity 'fingerprints', manifold learning and spatial statistics, to the resting-state fMRI data to obtain highly individualized representations of striatal functioning ('connectopic maps') for each subject (8). These maps represent slowly varying topographic patterns of connectivity ('connectopic gradients') that reveal how connectivity changes within a target region in relation to the rest of the brain. The striatum was defined as the target region based on the Harvard-Oxford Atlas and includes the putamen, nucleus accumbens and caudate nucleus. Although we focus on the principal or dominant gradient, multiple overlapping topographic representations can exist simultaneously within a single region, so both principal and second gradients were estimated to be able to investigate other potential effects driving associations to behavior. All gradients were visually inspected, and subjects were excluded if a clear gradient could not be estimated or spatial correlation of individual gradients to the group maps was low (n=17). A trend surface model was fit to the underlying gradient on the basis of the three-dimensional polynomial basis functions (8, 9). To determine the optimal model order, Bayesian Information Criterions (BIC) curves were computed across a range of model orders (1-12) at the group-level (9). The trend surface coefficients from each hemisphere were concatenated for each subject and entered into a penalized canonical correlation analysis (CCA) model, independently for the first and second gradients obtained (9). Using the functional gradients obtained, we also calculated the corresponding functional projections to the whole brain (8).

*CCA modelling and validation*
We used penalized CCA to determine the association between the behavioral domains of functioning and striatal gradients (Figure 1). This procedure has been described in detail in earlier publications (10). Briefly summarized; age and gender were regressed out of both behavioral and imaging data before entering them into the analysis. To ensure reproducibility, we embedded the CCA procedure in a split-half resampling procedure and also employed a stability selection approach (11) in order to reduce dimensionality in the brain space. Stability selection is a principled technique for variable selection that aims to select only the most stable features under repeated resamplings of the data and provides rigorous family wise error control over the selected variables (11). We stratified the data into a training and testing sample (50/50). Using the training data, we applied CCA to 100 subsamples (70% of the training data) and selected brain features that were consistently selected (top 20% range) across those 100 subsamples. The (L1) sparsity constraints were fixed at $\frac{1}{2}\sqrt{p}$, where $p$ is the number of features in each view, following the procedure outlined in Ing et al. (10).

Using the selected features, we performed the main CCA in the training data to obtain within-samples correlations between functional domains and striatal gradients ($\rho_{in}$). The canonical weights were then applied to the testing data and correlations were calculated with the new canonical correlates to obtain the out of sample correlation ($\rho_{out1}$). Then, the data were permuted (500 permutations) to obtain the corresponding null to compare against ($null_1$), using the same features since, as expected, permuted data did not result in any stable features being selected (see below). To reduce the risk of spurious results, the main CCA and subsequent steps were repeated after switching the training and testing sample. In doing so, we obtain another out of sample correlation $\rho_{out2}$ and another corresponding null ($null_2$). Note that feature selection was controlled using stability selection prior to permutation and was not embedded in the permutation procedure. We used a simple empirical test to ensure that feature selection did not introduce bias into the statistical testing procedure, by also repeating the stability selection procedure within the permutation test. After permuting the data, no features were deemed stable by the stability selection step, as expected. This indicates that the stability selection procedure was able to identify the truly informative features without overfitting to the data, which was expected on the basis of the theory underlying stability selection (11). We considered the model statistically significant when the mean of $\rho_{out1}$ and $\rho_{out2}$ was higher than 95% of the combined $null_1$ and $null_2$, corresponding to a significance level of $p < 0.05$. To establish model consistency, we performed the full analysis ten times using different (stratified) data splits. Finally, we calculated the (cross-)loadings of brain features to the different functional domains to explore the brain-behavior relationship in more detail to determine the amount of variance in the different functional domains that is explained by striatal connectopy.

**Additional analyses**

To determine other potential explanations for the observed interaction between the functional striatal gradients and the functional domains, the full analysis was repeated using three different data views: 1) classifications labels instead of functional domains, 2) striatal morphology (voxel-based morphometry) instead of striatal gradients, and 3) the second striatal gradients maps.

*1) classification labels*
Instead of the functional domains scores derived from the factor analysis, we used the classification labels for each of the four diagnostic categories included (major depressive disorder, anxiety disorder, attention deficit/hyperactivity disorder and/or autism spectrum disorder) into the CCA analysis. In the stability selection step, no stable brain features were observed. Using the full set of brain features (the trendsurface coefficients), the CCA model did not reach statistical significance (mean out-of-sample correlation 0.097 ; $p = 0.40$).

*2) striatal morphology*
Using the T1 images, the standard voxel-based morphometry pipeline from FSL (fsl.fmrib.ox.ac.uk/fsl/fslwiki/FSLVBM) was used to obtain VBM images of the striatum for each subject (2 844 voxels per subject). These were entered into the CCA with the functional domain scores. Stability selection on the brain data revealed no stable set of brain features across iterations. Using the full set of features, we found no significant association (mean out-of-sample correlation -0.06, $p = 0.51$).

*3) second striatal gradient maps*
Finally, instead of the primary gradients, we also checked for associations of the functional domain scores to the second connectopic maps obtained (see Figure S1). Again, within the CCA no stable features were obtained. Using the full set of gradient features, the CCA did not reveal a significant association between the functional domain scores and second gradient map coefficients (mean out-of-sample correlation = -0.038 ; $p = 0.36$).

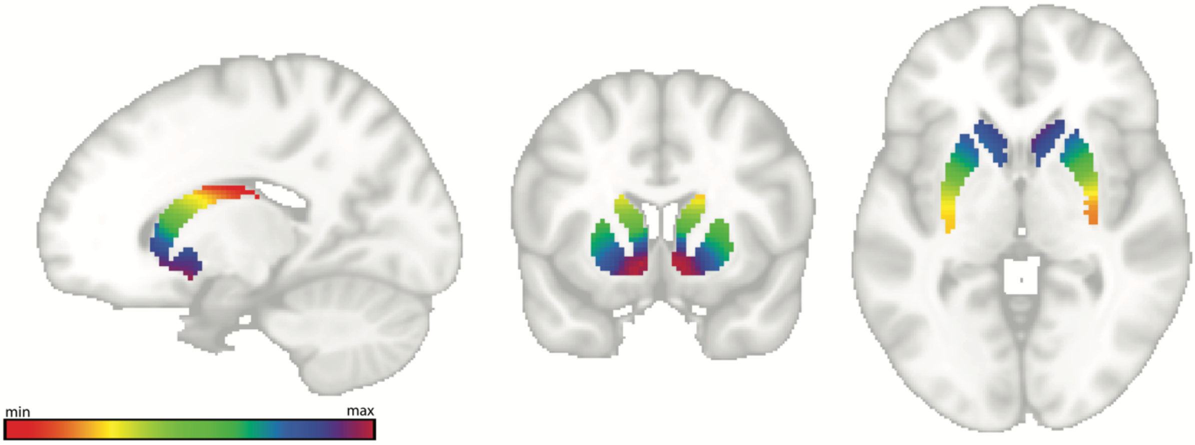


*Figure S1. Results of the second connectopic map*

**References**

1. van Eijndhoven PFP, Collard RM, Vrijsen JN, et al.: Measuring Integrated Novel Dimensions in Neurodevelopmental and Stress-related Mental Disorders (MIND-Set): a cross-sectional

comorbidity study from an RDoC perspective [Internet]. medRxiv 2021; 1–39[cited 2021 Nov 9] Available from: https://www.medrxiv.org/content/10.1101/2021.06.05.21256695v1
2. First M, Gibbon M, Spitzer R, et al.: Structured clinical interview for DSM-IV-TR axis I disorders, reseach version, patient edition (I/P). New York, New York: Biometrics Research, New York State Psychiatric Institute, 1997
3. Pettersson R, Söderström S, Nilsson KW: Diagnosing ADHD in Adults: An Examination of the Discriminative Validity of Neuropsychological Tests and Diagnostic Assessment Instruments. J Atten Disord 2018; 22:1019–1031
4. Vuijk R, Deen M, Arntz A, et al.: First Psychometric Properties of the Dutch Interview for Diagnostic Assessment of Autism Spectrum Disorder in Adult Males Without Intellectual Disability. J Autism Dev Disord 2021;
5. Horn JL: A rationale and test for the number of factors in factor analysis. Psychometrika 1965; 30:179–185
6. Jenkinson M, Beckmann CF, Behrens TEJ, et al.: FSL. Neuroimage 2012; 62:782–790
7. Pruim RHR, Mennes M, van Rooij D, et al.: ICA-AROMA: A robust ICA-based strategy for removing motion artifacts from fMRI data. Neuroimage 2015; 112:267–277
8. Haak K V., Marquand AF, Beckmann CF: Connectopic mapping with resting-state fMRI. Neuroimage 2018; 170:83–94
9. Marquand AF, Haak K V., Beckmann CF: Functional corticostriatal connection topographies predict goal-directed behaviour in humans. Nat Hum Behav 2017; 1
10. Ing A, Sämann PG, Chu C, et al.: Identification of neurobehavioural symptom groups based on shared brain mechanisms. Nat Hum Behav 2019 312 2019; 3:1306–1318
11. Meinshausen N, Bühlmann P: Stability selection. J R Stat Soc Ser B Stat Methodol 2010; 72:417–473
